# Does transcutaneous vagus nerve stimulation alter pupil dilation? A living Bayesian meta-analysis

**DOI:** 10.1101/2024.09.02.610851

**Authors:** Ipek Pervaz, Lilly Thurn, Cecilia Vezzani, Luisa Kaluza, Anne Kühnel, Nils B. Kroemer

## Abstract

Transcutaneous vagus nerve stimulation (tVNS) has emerged as a promising technique to modulate autonomic functions, and pupil dilation has been recognized as a promising biomarker for tVNS-induced monoaminergic release. Nevertheless, studies on the effectiveness of various tVNS protocols have produced heterogeneous results on pupil dilatation to date. Here, we synthesize the existing evidence and compare conventional continuous and pulsed stimulation protocols using Bayesian meta-analysis. To maintain a living version, we developed a Shiny App with the possibility to incorporate newly published studies in the future. Based on a systematic review, we included 18 studies (N = 771) applying either continuous or pulsed stimulation protocols. Across studies, we found anecdotal evidence for the alternative hypothesis that tVNS increases pupil size (*g* = 0.14, 95% CI = [0.001, 0.29], BF01 = 2.5). Separating studies according to continuous vs. pulsed protocols revealed that results were driven by studies using pulsed taVNS (strong evidence for the alternative hypothesis: *g* = 0.34, 95% CI = [0.15, 0.53], BF10 = 14.15) while continuous tVNS provided strong evidence for the null hypothesis (*g* = 0.01, CI = [-0.15, 0.16], BF01= 20.7). In conclusion, our meta-analysis highlights differential effects of continuous and pulsed tVNS protocols on pupil dilation. These findings underscore the relevance of tVNS protocols in optimizing its use for specific applications that may require modulation of tonic vs. phasic monoaminergic responses.

## Introduction

The vagus nerve is a crucial communication pathway between the brain and the body that innervates key visceral organs such as the heart, lungs, and stomach (Berthoud et al., 2021). By bidirectionally transmitting signals between the body and the brain, it helps maintaining bodily homeostasis (Chae et al., 2003; Breit et al., 2018) and is intricately involved in cognitive and affective processes such as motivation, reinforcement learning, and mood regulation (Berthoud and Neuhuber, 2000; Breit et al., 2018; Kühnel et al., 2020; Neuser et al., 2020; Ferstl et al., 2022; Teckentrup and Kroemer, 2023). Invasive vagus nerve stimulation (VNS) is approved as a treatment for epilepsy and depression (Johnson and Wilson, 2018) and has been explored for the treatment of various other disorders, including migraine, Parkinson’s and Alzheimer’s disease, anxiety, and eating disorders (Beekwilder and Beems, 2010; Wang et al., 2021). However, invasive VNS requires surgery with common associated risks and higher costs (Yang and Phi, 2019). Thus, developing non-invasive transcutaneous VNS (tVNS) protocols that provide safe and scalable use while robustly stimulating vagal afferent projections is essential for broader applications.

Non-invasive tVNS applies mild electrical stimulation to the skin in areas innervated by the vagus nerve. For stimulation of the auricular branch of the vagus nerve (taVNS), surface electrodes can be placed at the external ear, whereas electrodes on the neck stimulate the cervical branch of the vagus nerve (tcVNS) (Tekdemir et al., 1998; Kiyokawa et al., 2014; Yap et al., 2020). In general, tVNS is safe and well-tolerated (Redgrave et al., 2018). Moreover, it has shown comparable effectiveness as invasive VNS in the treatment of drug-resistant epilepsy (Bauer et al., 2016; Lampros et al., 2021) and major depressive disorder (Hein et al., 2013; Rong et al., 2016). Although tVNS is a promising alternative, several limitations halt its more widespread application in clinical settings. So far, conclusively large clinical trials are missing and recent small-scale studies have produced heterogeneous results across different behavioral and physiological outcomes. For example, Burger and colleagues have shown that tVNS accelerated fear extinction in one study (Burger et al., 2016), but did not replicate their findings in a later study (Burger et al., 2018). Such heterogeneity may partly be explained by anatomical differences (e.g., distance of vagal afferent fibers to the skin). However, except for tVNS-induced activation in the target region of vagal afferents in the brainstem, a reliable and scalable biomarker of an effective stimulation is missing (Burger et al., 2020b). Moreover, individual responses to tVNS may be influenced by age, gender, comorbidities, and medication use (Kemp et al., 2014; De Couck et al., 2017; Janner et al., 2018; Cakmak, 2019; Farmer et al., 2021). Other proposed biomarkers including heart rate variability (Wolf et al., 2021; Tarasenko et al., 2022), far-field potentials (Polak et al., 2009), and noradrenergic markers such as salivary alpha-amylase (Giraudier et al., 2022) and P300 have not demonstrated sufficiently robust effects of tVNS (for a review see (Burger et al., 2020b; Wolf et al., 2021). To conclude, there is a need for reliable biomarkers that would facilitate the development of optimized tVNS protocols tailored to individual characteristics and needs for the intended intervention.

Recently, tVNS-induced pupil dilation has gained traction as a suitable biomarker to track vagal afferent activation. Vagal afferent projections modulate the locus coeruleus-noradrenaline (LC-NA) system via projections from the nucleus of the solitary tract (NTS) in the brainstem (Adrienne and Guy, 2006; Dietrich et al., 2008; Hulsey et al., 2017; Yakunina et al., 2018). In turn, activation of the LC-NA system controls the pupil diameter (Mathôt, 2018). In rodents, VNS lead to increased LC activity (Hulsey et al., 2017) as well as pupil dilation (Bianca and Komisaruk, 2007; Mridha et al., 2021) in a dose-dependent manner with higher current intensities and longer pulse widths. In contrast to preclinical or invasive VNS studies, tVNS did not consistently elicit pupil dilation in humans (Burger et al., 2020b). Whereas several studies reported increased pupil dilation induced by tVNS (Capone et al., 2021; Omer et al., 2021; D’Agostini et al., 2023; Lloyd et al., 2023; Wienke et al., 2023), others did not obtain this result (Keute et al., 2019; Warren et al., 2019; Burger et al., 2020a; Borges et al., 2021; D’Agostini et al., 2021; D’Agostini et al., 2022; Villani et al., 2022; Anyla et al., 2023). This divergence might be explained by different tVNS protocols since the conventional protocol for clinical applications is a biphasic stimulation with 25 Hz (250 µs pulse width) for 30s ON/30s OFF (referred to as continuous stimulation). Even though continuous tVNS has been shown to robustly enhance BOLD responses in the NTS (Teckentrup et al., 2021), pupil dilation often remained unaffected (Warren et al., 2019; Burger et al., 2020a; Borges et al., 2021; D’Agostini et al., 2021; D’Agostini et al., 2022). In contrast, animal research typically uses shorter stimulation pulses ranging between 0.5–5 seconds (Mridha et al., 2021). Pulsed tVNS might more closely mimic the physiological cholinergic signals transmitted by the vagus nerve (Bowles et al., 2022), and thus might better drive the activity of the LC-NA system to induce pupil dilations (Reimer et al., 2016). Recent studies have used pulsed tVNS in humans as well and reported significant increases in pupil dilation (Sharon et al., 2021; D’Agostini et al., 2023; Lloyd et al., 2023; Wienke et al., 2023; Ludwig et al., 2024), but a systematic review and meta-analysis are missing so far.

To this end, we conducted a living Bayesian meta-analysis of studies comparing pupil dilation in response to tVNS. We further evaluated systematic differences between study protocols, such as the use of continuous or pulsed tVNS. Based on previous evidence, we hypothesized that tVNS leads to increased pupil phasic dilations when after short pulses of tVNS whereas continuous tVNS does not robustly increase the average pupil size. A robust increase in pupil dilation during tVNS would validate the potential of pupil dilation as a biomarker of vagus nerve activation. Moreover, if tVNS-induced pupil dilations are specific for pulsed stimulation protocols, this pave the way for to apply pulsed stimulation which more closely mimic physiological signals in settings where stimulation should be paired with specific events in a closed-loop setting.

## Methods

### Search strategies

The systematic literature search for tVNS and pupil responses was performed on 17/10/2023 in the databases Pubmed and Web of Science (keywords in the SI). In addition to the papers identified through the systematic database search, three papers were manually added after identification from preprint servers or notification by the authors. After screening abstracts, 20 papers were assessed. From these 20 papers, 3 papers had to be excluded (for details see SI), leaving 17 papers that were relevant for the meta-analysis (Figure 1). We extracted information from their figures using *WebPlotDigitizer* (Version 4.6, https://automeris.io/WebPlotDigitizer) when information was not provided and contacted authors from the papers that we could not obtain sufficient data from (Pandža et al., 2020; Konjusha et al., 2023). Finally, we included 15 papers reporting results from 18 studies (Table 1) in the meta-analysis. One paper included the results from 3 studies (Burger et al., 2020a) and for another paper we split up the results into two studies depending on the stimulation type continuous vs. pulsed to aid the comparison of the protocols (Skora et al., 2024). Missing studies with relevant results can be forwarded to the authors anytime and included to the Shiny App to incorporate any emerging evidence.

**Figure 1:**
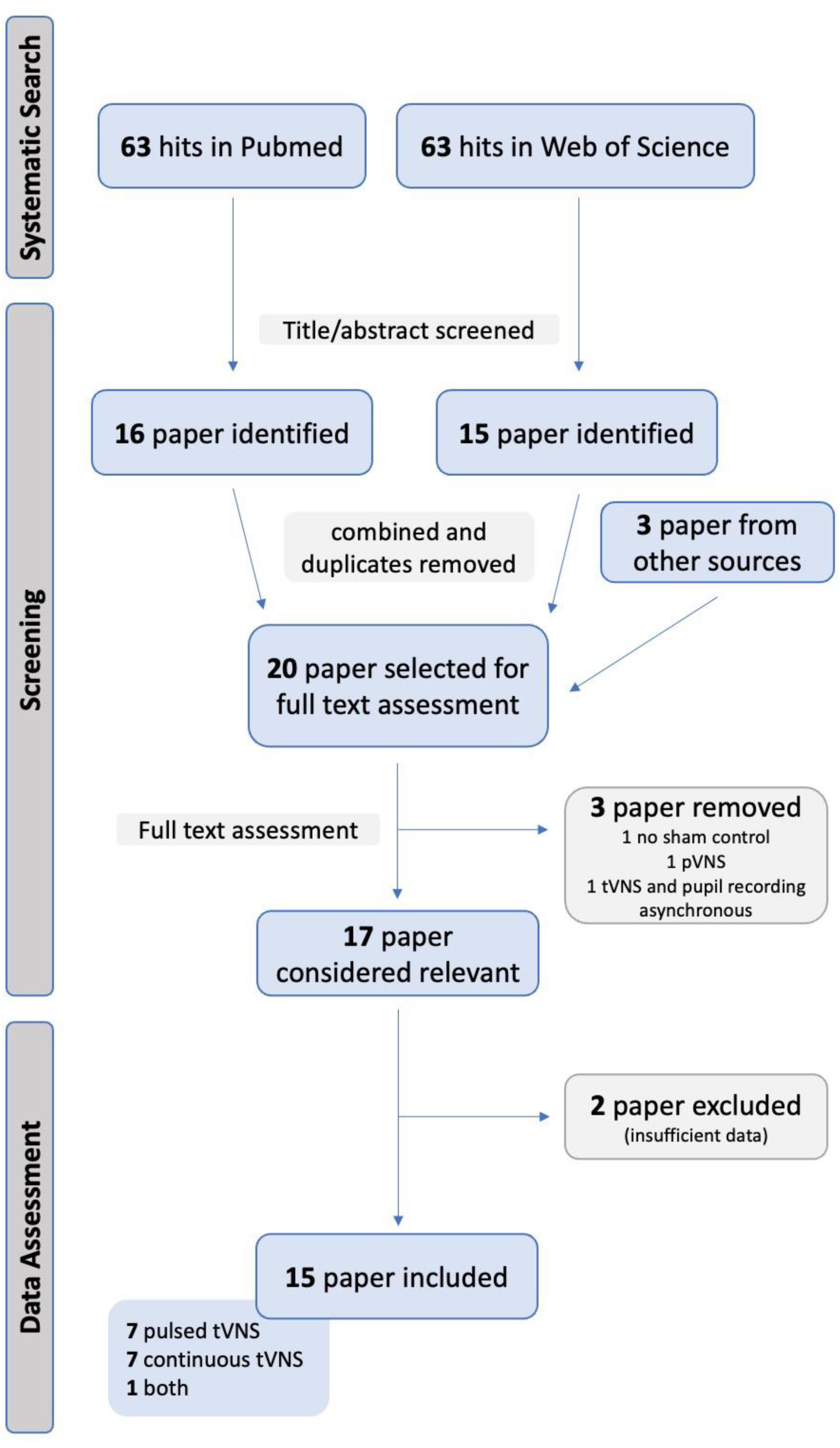
PRISMA flowchart showing the exclusion process for articles included in the meta- analysis

**Table 1:**
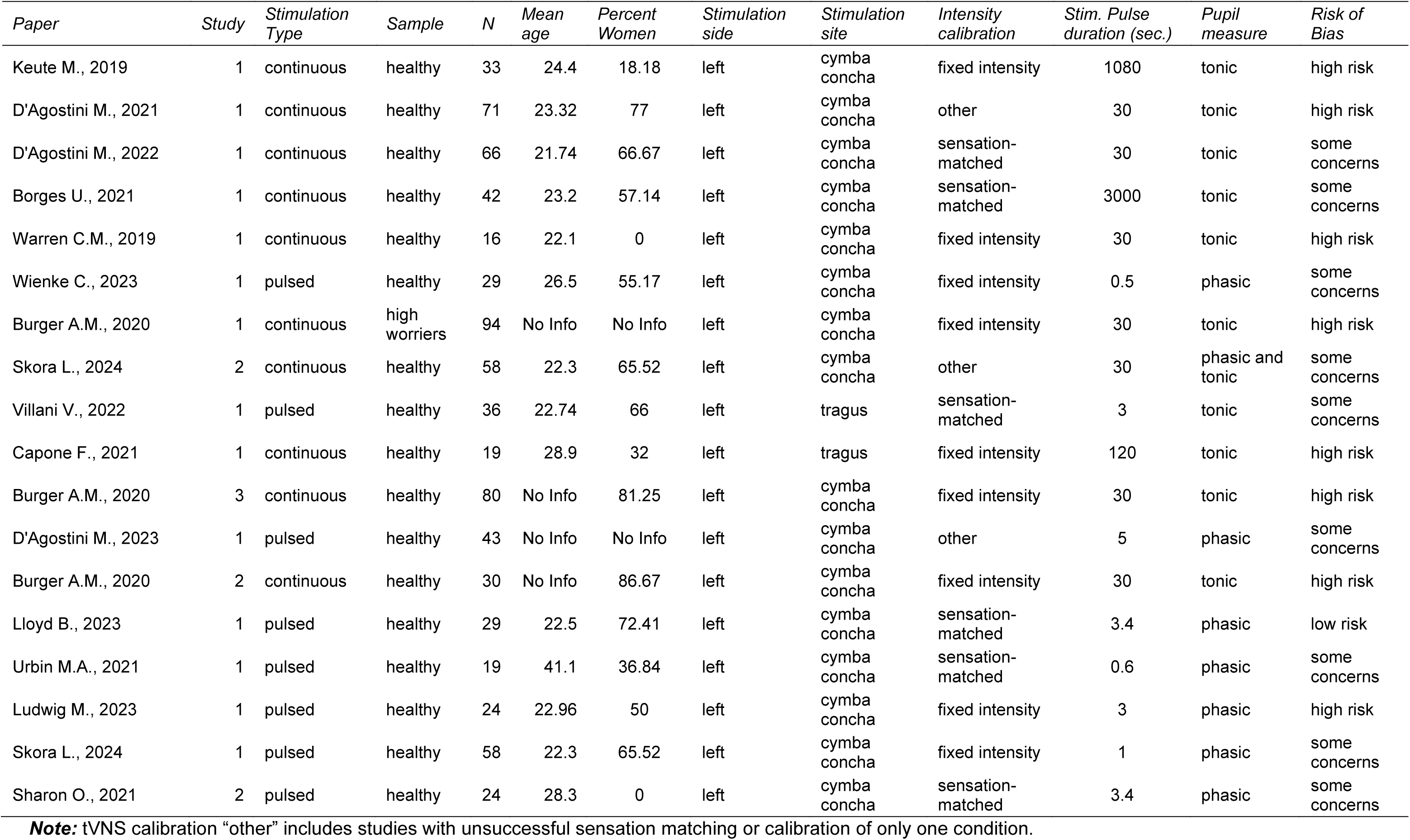
Studies included in the paper’s exemplary analysis. A more comprehensive table can be found in the Shiny app.

| *Paper* | *Study* | *Stimulation Type* | *Sample* | *N* | *Mean age* | *Percent Women* | *Stimulation side* | *Stimulation site* | *Intensity calibration* | *Stim. Pulse duration (sec.)* | *Pupil measure* | *Risk of Bias* |
| --- | --- | --- | --- | --- | --- | --- | --- | --- | --- | --- | --- | --- |
| Keute M., 2019 | 1 | continuous | healthy | 33 | 24.4 | 18.18 | left | cymba<br>concha | fixed intensity | 1080 | tonic | high risk |
| D'Agostini M., 2021 | 1 | continuous | healthy | 71 | 23.32 | 77 | left | cymba<br>concha | other | 30 | tonic | high risk |
| D'Agostini M., 2022 | 1 | continuous | healthy | 66 | 21.74 | 66.67 | left | cymba<br>concha | sensation-<br>matched | 30 | tonic | some<br>concerns |
| Borges U., 2021 | 1 | continuous | healthy | 42 | 23.2 | 57.14 | left | cymba<br>concha | sensation-<br>matched | 3000 | tonic | some<br>concerns |
| Warren C.M., 2019 | 1 | continuous | healthy | 16 | 22.1 | 0 | left | cymba<br>concha | fixed intensity | 30 | tonic | high risk |
| Wienke C., 2023 | 1 | pulsed | healthy | 29 | 26.5 | 55.17 | left | cymba<br>concha | fixed intensity | 0.5 | phasic | some<br>concerns |
| Burger A.M., 2020 | 1 | continuous | high<br>worriers | 94 | No Info | No Info | left | cymba<br>concha | fixed intensity | 30 | tonic | high risk |
| Skora L., 2024 | 2 | continuous | healthy | 58 | 22.3 | 65.52 | left | cymba<br>concha | other | 30 | phasic and<br>tonic | some<br>concerns |
| Villani V., 2022 | 1 | pulsed | healthy | 36 | 22.74 | 66 | left | tragus | sensation-<br>matched | 3 | tonic | some<br>concerns |
| Capone F., 2021 | 1 | continuous | healthy | 19 | 28.9 | 32 | left | tragus | fixed intensity | 120 | tonic | high risk |
| Burger A.M., 2020 | 3 | continuous | healthy | 80 | No Info | 81.25 | left | cymba<br>concha | fixed intensity | 30 | tonic | high risk |
| D'Agostini M., 2023 | 1 | pulsed | healthy | 43 | No Info | No Info | left | cymba<br>concha | other | 5 | phasic | some<br>concerns |
| Burger A.M., 2020 | 2 | continuous | healthy | 30 | No Info | 86.67 | left | cymba<br>concha | fixed intensity | 30 | tonic | high risk |
| Lloyd B., 2023 | 1 | pulsed | healthy | 29 | 22.5 | 72.41 | left | cymba<br>concha | sensation-<br>matched | 3.4 | phasic | low risk |
| Urbin M.A., 2021 | 1 | pulsed | healthy | 19 | 41.1 | 36.84 | left | cymba<br>concha | sensation-<br>matched | 0.6 | phasic | some<br>concerns |
| Ludwig M., 2023 | 1 | pulsed | healthy | 24 | 22.96 | 50 | left | cymba<br>concha | fixed intensity | 3 | phasic | high risk |
| Skora L., 2024 | 1 | pulsed | healthy | 58 | 22.3 | 65.52 | left | cymba<br>concha | fixed intensity | 1 | phasic | some<br>concerns |
| Sharon O., 2021 | 2 | pulsed | healthy | 24 | 28.3 | 0 | left | cymba<br>concha | sensation-<br>matched | 3.4 | phasic | some<br>concerns |

**Note:** tVNS calibration “other” includes studies with unsuccessful sensation matching or calibration of only one condition.

### taVNS stimulation protocols

taVNS stimulation is applied using various protocols with two broad categories emerging. First, in line with most clinical applications (Gerges et al.), the vagus nerve is stimulated continuously over several minutes to hours (for details SI). Second, inspired by animal research, the vagus nerve can be stimulated with a pulsed protocol that entails brief stimulation periods of a few seconds (0.6s (Urbin et al., 2021) - 5s (D’Agostini et al., 2023)) as would be expected by bodily feedback signals. For coding the studies, stimulation periods shorter or equal to 5s were considered as pulsed stimulation. Continuous stimulation protocols had at least 30s of uninterrupted stimulation.

In addition to the two stimulation protocol types, taVNS studies vary in the applied frequency, pulse width, and intensity (for more details see SI). One important consideration is the stimulation intensity, with 6 studies (2 continuous taVNS, 4 pulsed taVNS) determining it based on participants’ perceptual threshold (Urbin et al., 2021). Another 8 studies (6 continuous taVNS, 2 pulsed taVNS) used fixed intensity for all participants with no difference between the stimulation intensity applied during taVNS and sham. Last, four studies (2 continuous, 2 pulsed) used modified versions of the two approaches that did not ensure sensation-matching (for details see SI). However, no systematic differences between pulsed and continuous stimulation protocols were identified regarding the determination of stimulation intensity. Considering that any sensory or painful sensation increases alertness and affects pupil size (Wang and Munoz, 2015), it is crucial to evaluate whether stimulation intensities and associated subjective sensations differ systematically between stimulation protocols (Ludwig et al., 2024).

### Pupillometry

Pupil dilation serves as a promising biomarker for vagal afferent activation reflecting LC-NA noradrenergic activity (Reimer et al., 2016). Pupil dilation is measured using eye tracking and different measurement devices and settings were used across the included studies (for details see SI).

Depending on the type of taVNS stimulation (continuous vs pulsed), pupil dilation was often analyzed differently. For continuous tVNS, most papers analyzed changes in pupil size by averaging across the complete stimulation period therefore assessing tonic changes in pupil size. Alternatively, the tVNS effect on task-related pupil dilations is measured (for details see SI). For pulsed tVNS, pupil dilations are measured for short (a few seconds long) post-stimulation epochs and the maximal dilation per trial is determined in comparison to a pre-stimulus baseline comparable to the approach for task event-related pupil dilations. Skora and colleagues also used this event-related phasic analysis approach for their continuous taVNS stimulation (Skora et al., 2024). They analyzed pupil response in the short epochs (∼5s) starting from the stimulation onset of each 30s ON phase and baseline-corrected using the last seconds of the previous 30s OFF period (Skora et al., 2024).

### Inclusion and exclusion criteria

To be included in the analysis, studies had to use tVNS and include sham stimulation as a control condition. However, we only found studies using auricular tVNS, therefore we only refer to taVNS in the results section. Furthermore, studies had to measure pupil size and report either pupil size or pupil dilation during stimulation and a few seconds after stimulation for the phasic responses. Only primary studies in humans were included and studies in animals were excluded during the screening process.

### Coding procedure

For all screened papers, we (IP and LT) coded those characteristics (for a full list, see SI). For studies not reporting the mean and standard deviation (SD), missing values were calculated and if the standard error (SE) was reported, SD was calculated using *SD = SE * sqrt(Ngroup).* For studies reporting only the median and interquartile ranges (IQR) we used the *metafor* package (Viechtbauer, 2010) to estimate the mean and SD. From the resulting data sets, effect sizes (Hedges’ *g*) were calculated using the same *metafor* package in R version 4.2.1. Finally, for the studies reporting pupil size before and during stimulation rather than pupil dilation (Warren et al., 2019; Burger et al., 2020a; Capone et al., 2021; D’Agostini et al., 2021; D’Agostini et al., 2022), we calculated baseline-corrected pupil sizes before estimating the effect size.

### Risk of bias

The risk of bias was assessed using the open-access revised Cochrane risk of bias tool for randomized trials *(RoB2*, (Sterne et al., 2019)). Two of the authors (IP and LK) conducted the assessment independently. Differences in the risk of bias assessment were then reviewed by a third author (AK). The studies were categorized as low risk, medium risk, or high risk in five different domains by evaluating the randomization quality, the blinding, the effect of potential missing data, the validity of the method used to measure the outcome, and whether the study was conducted following a pre-registered analysis plan. We classified studies with a high risk or medium risk of bias in more than one domain as studies with a high risk of bias. If only one domain was classified as high or medium risk of bias, we classified the study as having a medium risk of bias (for detailed coding of the domains, see SI). Overall (Figure 2D), only one study (Lloyd et al., 2023) was categorized as low risk for bias, eight studies as having some risk (Borges et al., 2021; Sharon et al., 2021; Urbin et al., 2021; D’Agostini et al., 2022; Villani et al., 2022; D’Agostini et al., 2023; Wienke et al., 2023; Skora et al., 2024), and eight studies (Keute et al., 2019; Warren et al., 2019; Burger et al., 2020a; Capone et al., 2021; D’Agostini et al., 2021; Ludwig et al., 2024) were categorized as high risk of bias.

**Figure 2:**
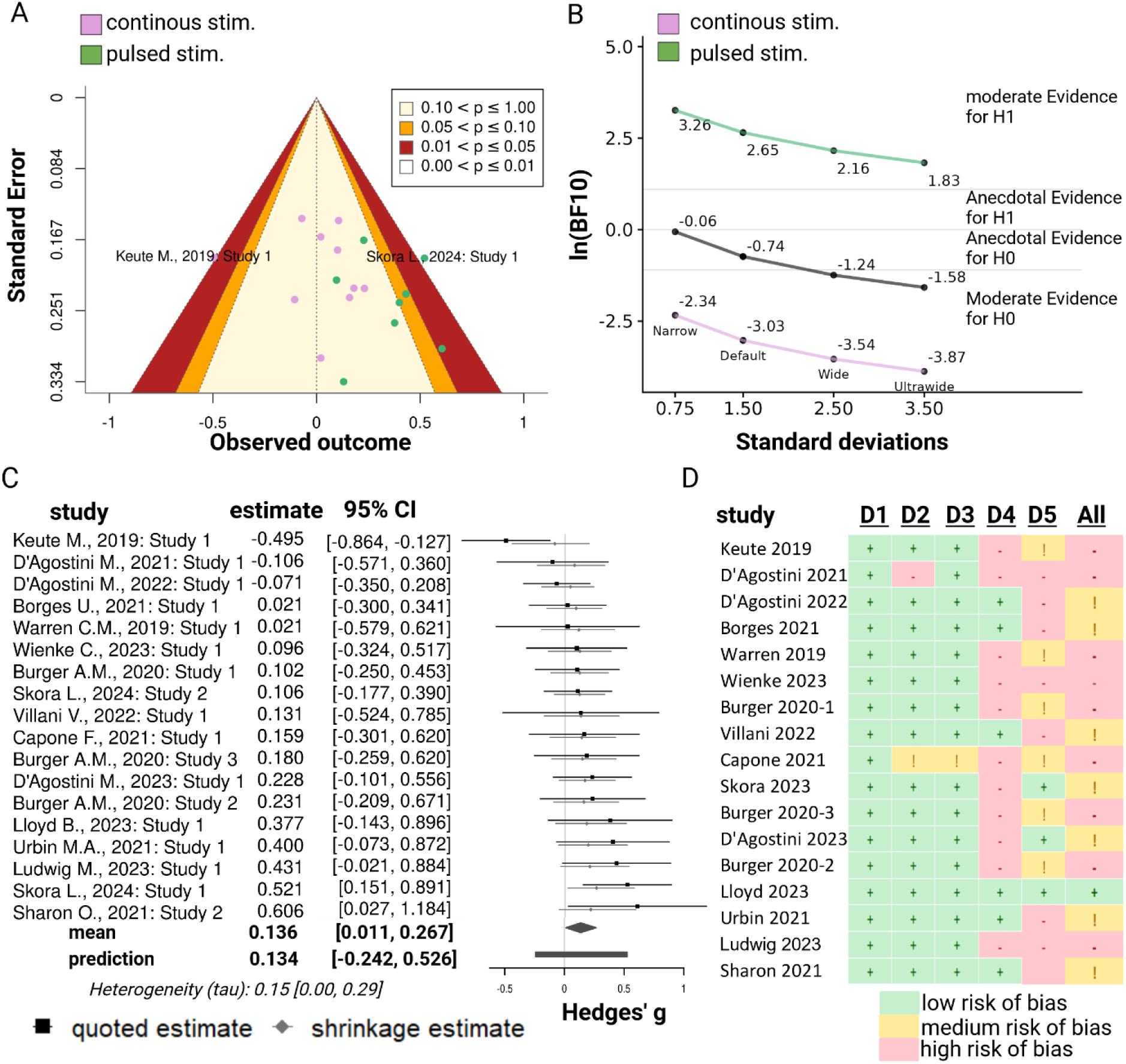
Across 17 studies, the meta-analysis shows moderate evidence against an effect of taVNS on pupil dilation. **A**: Contour-enhanced funnel plot showing standard error and observed outcome for each included study, grouped by stimulation type. Studies applying continuous (tonic) stimulation are depicted in pink, while studies applying pulsed (phasic) stimulation are depicted in green. **B**: Bayes factor (BF, log-transformed) robustness plot. BF are plotted across prior distributions differing in their width. The plot shows moderate evidence for H0 across all studies (in black). After grouping by stimulation type, we find strong evidence for H0 for continuous stimulation (in pink) and moderate evidence for H1 for pulsed stimulation (in green). **C**: Forest plot of all included studies, showing a no effect across all included studies (mean = 0.14, CI [0.01, 0.27] for taVNS compared to sham. D: We assessed the risk of bias for all studies using a standardized checklist (Sterne et al., 2019). Studies are rated across 5 domains (D1: Randomization D2: Deviations from the intervention D3: Missing data D4: Validity of the outcome measure D5: Preregistration). If a study showed increased risk, we rated them as medium risk across all domains and as high risk if more than one domain showed an increase risk of bias. Only one study (Lloyd et al., 2023) was rated to have a low risk of bias.

### Design of the Shiny app

To build the Shiny app, we adapted a previously developed app by our group (Wolf et al., 2021). Analogous to the previous living meta-analysis using taVNS-induced changes in heart rate variability as the outcome, the Shiny app allows the user to change the inclusion criteria for the selection of studies into the meta-analysis (example choices in the SI). After selecting the desired parameters and pressing the *Re-Calculate Meta-Analysis* button, the app will perform the corresponding analysis. Since most papers contain multiple effect sizes, narrowing the inclusion parameters in the Shiny App may result in changes in the reported effect sizes of these papers. The app provides a *Forest plot* of all included studies (Figure 1), a *Funnel plot* to assess publication bias (Figure 2) as well as outliers check, and plots of prior and posterior densities for heterogeneity (τ) and effect (μ). R Code for the Shiny app is available on GitHub (https://github.com/neuromadlab/taVNSpupilmeta). The complete analyses and inclusion criteria can be found in the Shiny app accompanying the paper (https://neuromadlab.shinyapps.io/tVNSPupilmeta/).

## Statistical methods

As a measure of effect size, we calculated *Hegdes’ g* (Hedges and Olkin, 1984) for each comparison between taVNS and sham based on means and SDs. To this end, we used the *metafor* package’s *escalc* function in R 4.2.1 (Viechtbauer, 2010). We performed a Bayesian random-effects meta-analysis using the *bayesmeta* package (Röver, 2020). Bayesian random-effects analyses are advantageous when the number of studies is still limited and they allow quantification of evidence in support of the null hypothesis in addition to testing the alternative hypothesis (Röver, 2020; Harrer et al., 2021; Wolf et al., 2021). Moreover, a Bayesian meta-analysis allows us to plot prediction intervals to make inferences about future studies not yet included in the meta-analysis (Higgins et al., 2008; Wolf et al., 2021). To compare taVNS-induced pupil responses between phasic vs. continuous stimulation protocols, we performed a Bayesian meta-regression with one factor using the *bmr* function of the *bayesmeta* package (Röver and Friede, 2023)

In line with previous work, we used a weakly informative prior with mean=0 and SD=1.5 for μ, and an informative half-Cauchy prior with a 0.5 scale for τ as default settings (Röver, 2020; Wolf et al., 2021). The Shiny app allows for the selection of different μ and τ priors, although such changes must be made with caution (van Doorn et al., 2021). Here, we report Bayes factors (BF) and shortest credible intervals for the difference in pupil dilation between taVNS and sham stimulation, as well as for the heterogeneity parameter (Figure 2B; (Liu et al., 2015; Wolf et al., 2021)). The Shiny app automatically reports the BF calculated using the SD selected by the user (“default”), as well as an “ultrawide” (SD+2), “wide” (SD+1), and “narrow” (SD/2) BF (Figure 2B).

## Results

To evaluate if taVNS exerts robust effects on pupil dilation, we first compared the taVNS vs. sham stimulation using Bayesian random-effects meta-analysis. Based on our hypothesis, we investigated if taVNS-induced changes in pupil dilation depend on the use of continuous or pulsed stimulation protocols. Following recommendations (Allbritton et al., 2024), we provide a Shiny App to enable users to manipulate the analysis specifications and further explore the available data without relying on narrow inclusion decisions. To this end, we included 15 papers, containing 18 studies.

Across all studies using taVNS, the meta-analysis showed moderate evidence against a taVNS-induced increase in pupil size (*g* = 0.11, 95%CI = [-0.01,0.24], BF01=4.05 Figure 2), with significant evidence of heterogeneity among studies (τ= 0.12, 95%CI= [0.00,0.27], Figure 2C). The Bayes’ factor robustness analysis revealed that evidence against an effect of taVNS on pupil dilation was moderate for the default prior width and increased with increasing prior widths (Figure 2B). An outlier check revealed one potential outliers (Keute et al., 2019) with effect sizes of *g*= -0.49, 95%CI= [-0.86,- 0.13]. If this study was excluded from the analysis (i.e., comparable to a trimmed mean), it resulted in weak evidence for the alternative hypothesis (*g*= 0.12, 95%CI= [0.03,0.25], BF10= 2.08). The funnel plot showed weak skewness, indicating some degree (Figure 2A).

## Pulsed but not continuous taVNS increases pupil dilation

To determine whether the effects of taVNS on pupil dilation depend on the stimulation protocol, we performed separate meta-analyses for studies using continuous or pulsed stimulation protocols. In studies using continuous stimulation, taVNS did not significantly increase pupil dilation with strong evidence for the null hypothesis (*g* = 0.01, 95%CI = [-0.15,0.16], BF01= 20.7, Figure 3A). Evidence for the null hypothesis was consistent across prior widths (Figure 2B). The studies using continuous taVNS showed significant heterogeneity (τ= 0.11, 95%CI = [0.00,0.29]), and one study was identified as a potential outlier (Keute et al., 2019) with the effect size of *g* = -0.49, 95%CI = [-0.86,-0.13]. However, if this study was excluded, there was only anecdotal evidence against the null hypothesis (*g*= 0.06, 95%CI = [-0.08, 0.21], BF01= 0.93) indicating that more evidence would be necessary. The funnel plot showed no discernible skewness among the studies using continuous taVNS and thus no clear evidence of publication bias (Figure 2A).

**Figure 3.**
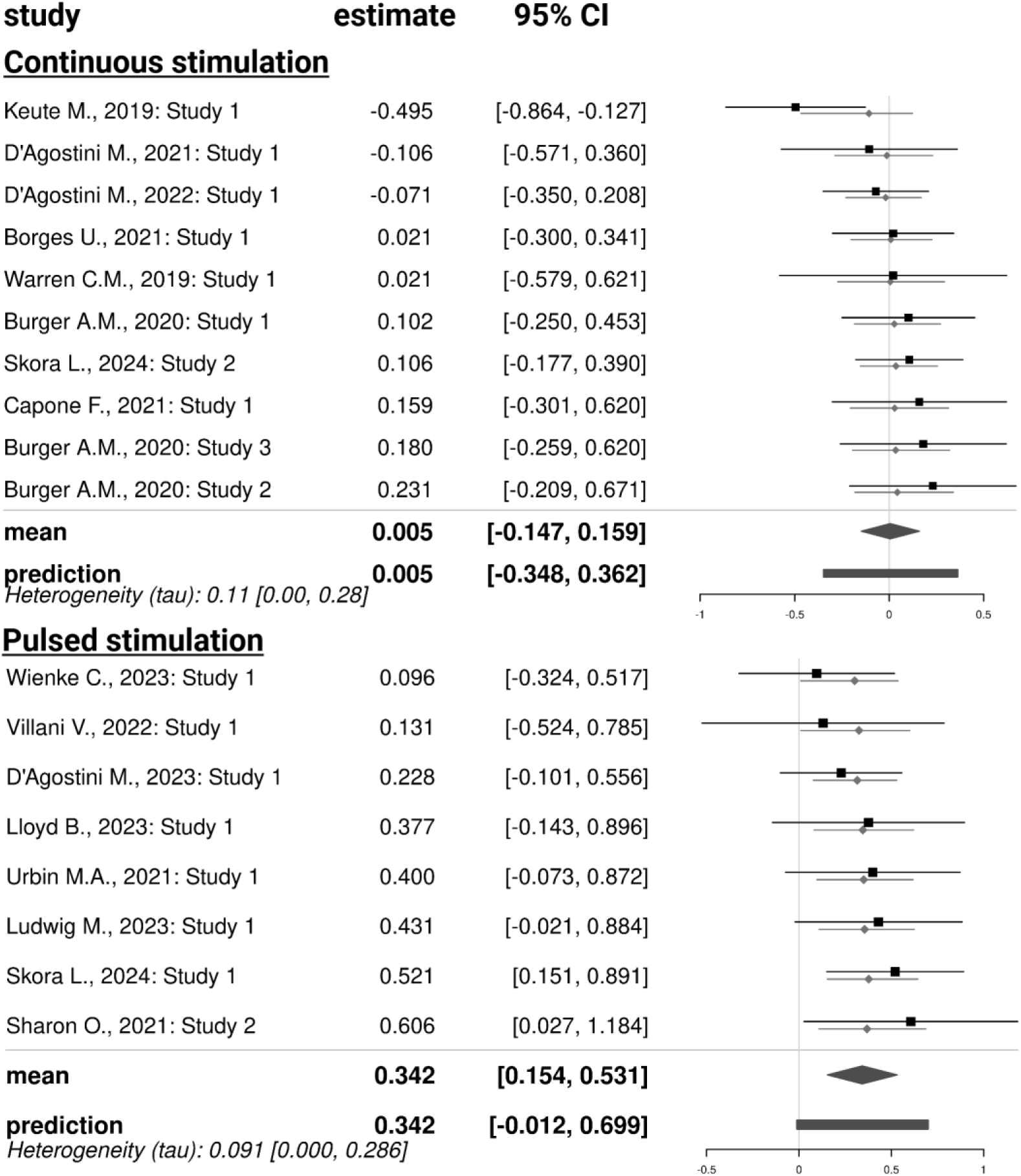
Stimulation effects of taVNS on pupil dilation depend on the use of a phasic vs. continuous protocol. Forest plot of all studies using continuous taVNS (upper) and pulsed taVNS (lower) with the effect size Hedge’s *g* and 95% CI, showing no effect of taVNS on pupil dilation under continuous stimulation (*g* = 0.01, 95% CI = [-0.15, 0.16], BF01 = 20.7), and strong evidence under pulsed stimulation (*g* = 0.34, 95% CI = [0.15, 0.53], BF10 = 14.15).

In studies using pulsed stimulation, taVNS increased pupil dilation with strong supporting evidence (*g*= 0.34, 95%CI = [0.15, 0.53], BF10= 14.15, Figure 3B). Although none of the studies was identified as an outlier, the funnel plot indicated some degree of publication bias (Figure 2A). Crucially, comparing the evidence for pulsed vs. continuous taVNS protocols using meta-regression showed that pulsed taVNS did induce a significantly higher pupil response than continuous stimulation (*g*pulsed – *g*continuous = 0.34, 95%CI = [0.11,0.56]).

## Risk of bias assessment

To evaluate whether our results are driven by studies with a high risk of bias, we performed the same analyses after excluding 8 studies classified as high risk. Across both stimulation protocols, the results showed anecdotal evidence for an increase in pupil size with taVNS compared to sham (g= 0.15, 95%CI= [-0.03,0.35], BF01= 1.3) indicating that more evidence from high-quality studies is necessary. When excluding high-risk studies from the meta-analysis of pulsed stimulation, the results still showed strong evidence for a taVNS-induced increase in pupil size (*g* = 0.36, 95%CI = [0.11,0.61], BF10 = 32.8).

## Discussion

tVNS has received increasing attention due to its easy convenient application and promising behavioral effects. However, a reliable and scalable biomarker to verify effective taVNS is still lacking. As one of the promising candidates, pupil dilation has produced heterogeneous results in response to taVNS so far. To address the heterogeneity among studies in the quickly growing field, we conducted a living Bayesian meta-analysis that examines studies measuring pupil dilation in response to taVNS (vs. sham). Crucially, we compared the conventional continuous stimulation with pulsed stimulation protocols that are often used in preclinical studies. Across all studies we found a small increase in pupil dilation but only with anecdotal evidence. However, comparing pulsed and continuous stimulation protocols revealed that only pulsed stimulation leads to a phasic increase in pupil diameter, whereas continuous taVNS does not increase tonic pupil size. To summarize, our results suggest that taVNS induces a phasic pupil response indicative of transient activation of the LC via vagal afferents. In contrast, tVNS does not lead to tonic increases in pupil dilation, suggesting that clinical and behavioral changes in settings with longer stimulation periods are not explained by tonic changes in LC signaling. Taken together, our results highlight the potential of novel pulsed stimulation protocols that are also common in preclinical research to induce phasic LC activity. Ultimately, such stimulation pulses might be combined with specific events or behaviors in a closed-loop setting to modify learning and decision-making.

Across all studies, taVNS increased pupil dilation compared to sham. Such taVNS-induced pupil dilations suggest effective activation of the vagus nerve and its downstream regions. The NTS in the brainstem is the first projection target of vagal afferent fibers (Frangos et al., 2015; Yakunina et al., 2017; Butt et al., 2020) and reliably activated by continuous taVNS (Frangos et al., 2015; Teckentrup et al., 2021). In turn, activation of the vagal afferent pathway also activates the LC-NA system as indexed by pupillary responses (Yakunina et al., 2017; Mathôt, 2018). Further exploration of stimulation protocols used across studies revealed that the increase in pupil size was driven by pulsed taVNS. This result is consistent with research in rodents, showing pupil dilation and cholinergic activation after pulsed (∼1-10s) stimulation of the vagus nerve (Collins et al., 2021; Mridha et al., 2021). Stronger phasic responses might indicate that pulsed taVNS better mimics natural phasic signaling of the vagus nerve that drives phasic activation of the LC-NA system (Reimer et al., 2016; Bowles et al., 2022). Research in animals has shown that rapid increases in pupil size reflect the phasic activity of NA-releasing axons (Reimer et al., 2016). Remarkably, continuous taVNS also resulted in phasic dilations of the pupil in one study that included both conditions (Skora et al., 2024). Phasic monoaminergic signals from the LC play a crucial role in encoding unsigned prediction errors or “surprise” signals (Dayan and Yu, 2006; Preuschoff et al., 2011). They are essential for learning as they help adjust future behavior by recalibrating expectations based on discrepancies between anticipated and actual outcomes (Deng et al., 2023). By enhancing phasic monoaminergic responses, taVNS might improve learning processes supporting more effective learning and conditioning, including during extinction learning (Burger et al., 2016; Burger et al., 2018; Szeska et al., 2020) or motor learning as previously shown in rodents (Bowles et al., 2022). Such event-locked interventions could be used to treat various mental disorders with disrupted learning processes, such as mood and anxiety disorders (Pike and Robinson, 2022). To conclude, the synthesized effects of pulsed taVNS on pupil dilation as a marker of phasic noradrenaline release spark hope for future clinical applications that improve aberrant learning.

Although the meta-analysis provides strong evidence for taVNS-induced pupil dilations using pulsed stimulation protocols, continuous taVNS did not elicit robust (tonic) increases in pupil dilation and there are several possible explanations for this. First, continuous taVNS may recruit differential responses over time that do not necessarily activate the LC-NA system for longer than an initial burst. This interpretation is supported by the study of Skora et al. (Skora et al., 2024). Other studies that averaged pupil dilation responses across longer periods might miss such a phasic response in their analysis (Skora et al., 2024). Second, phasic pupil dilation may primarily reflect salience-driven responses, whereas prolonged responses (e.g., to continuous taVNS) might reflect a longer-lasting stimulation of the LC-NA system via vagal afferents (Nieuwenhuis et al., 2011; Vespa et al., 2022). Even sham stimulation induces phasic pupil dilations that correspond to orienting responses to the sensory stimulus (Omer et al., 2021; Urbin et al., 2021). To minimize such confounds, we only included studies with suitable sham control of sensory effects in the meta-analysis. If the sensory experience is matched between conditions, taVNS-induced pupil dilations indicate that such responses are not merely due to the sensory aspect of the stimulation. Crucially, ∼50% of the pulsed stimulation studies used a sensation- matching method, whereas only ∼30% of the continuous stimulation studies did so, indicating that this aspect is unlikely to explain larger effects of pulsed stimulation. Third, even if continuous taVNS does not robustly alter pupil dilation, it might modulate noradrenaline on a different timescale (e.g., stimulation across hours or days, (Giraudier et al., 2022)). At a conceptual level, behavioral effects of continuous taVNS on mood and depression (Rong et al., 2016; Ferstl et al., 2022) or motivation and learning (Neuser et al., 2020; Szeska et al., 2020; Weber et al., 2021; Ferstl et al., 2024), indicate that such interventional effects are not necessarily dependent on driving phasic noradrenergic release.

Despite the strengths of a living Bayesian meta-analysis in emerging fields (Wolf et al., 2021), several limitations remain to be addressed in future work. First, the number of studies is low and the power of each study is limited. Future studies should therefore enroll more participants to improve the robustness of the reported findings. For example, with an expected small to medium effect size (d = .35) one would have to include 102 participants per group in a between-subject design to test for an increase in pupil size with 80% power. The power can be increased by within-subject designs depending on the reliability of the pupil response. Given the interactive nature of the Shiny App, future studies can be easily added to update the conclusions based on the advances in the field. Second, several studies showed a high risk of bias, mainly due to insufficient sensation matching between stimulation conditions and the absence of preregistration. Crucially, future studies should transparently report on preregistered outcomes (based on a detailed analysis plan) to reduce the risk of biases. Third, most studies differ in terms of their stimulation protocols (e.g., stimulation intensity, amplitude, and frequency) as well as pupil-related outcomes (e.g., task-related ERPDs, tonic increases, left or right or both eyes). Synthesis across such outcomes can be reconsidered with more available studies. Several animal studies and human studies have shown that stimulation parameters influence the size of the effect of taVNS on pupil size (Hulsey et al., 2017; Mridha et al., 2021; D’Agostini et al., 2023; Ludwig et al., 2024). Notably, only one study has directly compared continuous and pulsed taVNS (Skora et al., 2024) and all included studies used left auricular tVNS. Since lateralized effects have been shown for other outcomes (De Couck et al., 2017; Neuser et al., 2020; Teckentrup et al., 2020) and cervical tVNS migh recruit different afferents (Farmer et al., 2021; Jigo et al., 2024), future studies should focus on such gaps to improve the generalization to potential interventions.

Finding a reliable and scalable biomarker for the successful stimulation of the vagus nerve via tVNS is an important step for real-life applications. Our living Bayesian meta-analysis offers a robust statistical framework to assess the impact of taVNS on pupil dilation. Our results indicate that pulsed taVNS increases pupil dilation while continuous taVNS does not robustly alter tonic pupil dilation. Based on this systematic review of the available evidence to date, our study highlights the potential of pulsed taVNS to modify behavior that is dependent on eliciting phasic monoaminergic responses such as learning. Although heterogeneous study designs raise challenges for evidence synthesis, they also provide an opportunity to explore the most effective stimulation protocols across (independent) studies and teams. Crucially, the open format of our approach supports updating of the results as new studies become available and meta-regression may help identify optimized settings for future applications. In the long run, the curation of living meta-analysis on crucial outcomes of neuromodulation interventions could help improve targeted (e.g., specific groups or situations) interventions for novel applications.

## Acknowledgement

We thank Lisa Steffens for her support in coding the Shiny App. The study was supported by the BONFOR Grant O-1282.1 and the German Research Foundation (DFG), grant KR 4555/10-1.

## Author contributions

AK and NBK were responsible for the concept and design of the meta-analysis. IP and LT performed the literature search and coded eligible studies, and AK confirmed the coding. IP, LK, and AK performed the risk of bias assessment. CV performed the data analysis and AK, LT, & NBK contributed to analyses. IP, LT, AK, & NBK wrote the manuscript. All authors contributed to the interpretation of findings, provided critical revision of the manuscript for important intellectual content, and approved the final version for publication.

## Financial disclosure

The authors declare no competing financial interests.

## Notes

### Competing Interest Statement

The authors have declared no competing interest.

## References

1. Adrienne ED, Guy D (2006) Effect of Vagus Nerve Stimulation on Serotonergic and Noradrenergic Transmission. Journal of Pharmacology and Experimental Therapeutics 318:890.

2. Allbritton D, Gómez P, Angele B, Vasilev M, Perea M (2024) Breathing Life Into Meta-Analytic Methods. Journal of Cognition.

3. Anyla K, Shijing Y, Moritz M, Lorenza C, Tjalf Z, Christian B (2023) Auricular Transcutaneous Vagus Nerve Stimulation Specifically Enhances Working Memory Gate Closing Mechanism: A System Neurophysiological Study. The Journal of Neuroscience 43:4709.

4. Bauer S, Baier H, Baumgartner C, Bohlmann K, Fauser S, Graf W, Hillenbrand B, Hirsch M, Last C, Lerche H, Mayer T, Schulze-Bonhage A, Steinhoff BJ, Weber Y, Hartlep A, Rosenow F, Hamer HM (2016) Transcutaneous Vagus Nerve Stimulation (tVNS) for Treatment of Drug-Resistant Epilepsy: A Randomized, Double-Blind Clinical Trial (cMPsE02). Brain Stimul 9:356–363.

5. Beekwilder JP, Beems T (2010) Overview of the clinical applications of vagus nerve stimulation. J Clin Neurophysiol 27:130–138.

6. Berthoud H-R, Neuhuber WL (2000) Functional and chemical anatomy of the afferent vagal system. Autonomic Neuroscience 85:1–17.

7. Berthoud H-R, Albaugh VL, Neuhuber WL (2021) Gut-brain communication and obesity: understanding functions of the vagus nerve. The Journal of Clinical Investigation 131.

8. Bianca R, Komisaruk BR (2007) Pupil dilatation in response to vagal afferent electrical stimulation is mediated by inhibition of parasympathetic outflow in the rat. Brain Research 1177:29–36.

9. Borges U, Pfannenstiel M, Tsukahara J, Laborde S, Klatt S, Raab M (2021) Transcutaneous vagus nerve stimulation via tragus or cymba conchae: Are its psychophysiological effects dependent on the stimulation area? International Journal of Psychophysiology 161:64–75.

10. Bowles S, Hickman J, Peng X, Williamson WR, Huang R, Washington K, Donegan D, Welle CG (2022) Vagus nerve stimulation drives selective circuit modulation through cholinergic reinforcement. Neuron 110:2867–2885.e2867.

11. Breit S, Kupferberg A, Rogler G, Hasler G (2018) Vagus Nerve as Modulator of the Brain–Gut Axis in Psychiatric and Inflammatory Disorders. Frontiers in Psychiatry 9.

12. Burger AM, Van der Does W, Brosschot JF, Verkuil B (2020a) From ear to eye? No effect of transcutaneous vagus nerve stimulation on human pupil dilation: A report of three studies. Biological Psychology 152:107863.

13. Burger AM, D’Agostini M, Verkuil B, Van Diest I (2020b) Moving beyond belief: A narrative review of potential biomarkers for transcutaneous vagus nerve stimulation. Psychophysiology 57:e13571.

14. Burger AM, Verkuil B, Van Diest I, Van der Does W, Thayer JF, Brosschot JF (2016) The effects of transcutaneous vagus nerve stimulation on conditioned fear extinction in humans. Neurobiol Learn Mem 132:49–56.

15. Burger AM, Van Diest I, van der Does W, Hysaj M, Thayer JF, Brosschot JF, Verkuil B (2018) Transcutaneous vagus nerve stimulation and extinction of prepared fear: A conceptual non-replication. Sci Rep 8:11471.

16. Butt MF, Albusoda A, Farmer AD, Aziz Q (2020) The anatomical basis for transcutaneous auricular vagus nerve stimulation. J Anat 236:588–611.

17. Cakmak YO (2019) Concerning Auricular Vagal Nerve Stimulation: Occult Neural Networks. Frontiers in Human Neuroscience 13.

18. Capone F, Motolese F, Di Zazzo A, Antonini M, Magliozzi A, Rossi M, Marano M, Pilato F, Musumeci G, Coassin M, Di Lazzaro V (2021) The effects of transcutaneous auricular vagal nerve stimulation on pupil size. Clinical Neurophysiology 132:1859–1865.

19. Chae JH, Nahas Z, Lomarev M, Denslow S, Lorberbaum JP, Bohning DE, George MS (2003) A review of functional neuroimaging studies of vagus nerve stimulation (VNS). J Psychiatr Res 37:443–455.

20. Collins L, Boddington L, Steffan PJ, McCormick D (2021) Vagus nerve stimulation induces widespread cortical and behavioral activation. Curr Biol 31:2088–2098.e2083.

21. D’Agostini M, Burger AM, Villca Ponce G, Claes S, von Leupoldt A, Van Diest I (2022) No evidence for a modulating effect of continuous transcutaneous auricular vagus nerve stimulation on markers of noradrenergic activity. Psychophysiology 59:e13984.

22. D’Agostini M, Burger AM, Franssen M, Perkovic A, Claes S, von Leupoldt A, Murphy PR, Van Diest I (2023) Short bursts of transcutaneous auricular vagus nerve stimulation enhance evoked pupil dilation as a function of stimulation parameters. Cortex 159:233–253.

23. D’Agostini M, Burger AM, Franssen M, Claes N, Weymar M, von Leupoldt A, Van Diest I (2021) Effects of transcutaneous auricular vagus nerve stimulation on reversal learning, tonic pupil size, salivary alpha-amylase, and cortisol. Psychophysiology 58:e13885.

24. Dayan P, Yu AJ (2006) Phasic norepinephrine: a neural interrupt signal for unexpected events. Network 17:335–350.

25. De Couck M, Cserjesi R, Caers R, Zijlstra WP, Widjaja D, Wolf N, Luminet O, Ellrich J, Gidron Y (2017) Effects of short and prolonged transcutaneous vagus nerve stimulation on heart rate variability in healthy subjects. Autonomic Neuroscience 203:88–96.

26. Deng Y, Song D, Ni J, Qing H, Quan Z (2023) Reward prediction error in learning-related behaviors. Frontiers in Neuroscience 17.

27. Dietrich S, Smith J, Scherzinger C, Hofmann-Preiß K, Freitag T, Eisenkolb A, Ringler R (2008) A novel transcutaneous vagus nerve stimulation leads to brainstem and cerebral activations measured by functional MRI / Funktionelle Magnetresonanztomographie zeigt Aktivierungen des Hirnstamms und weiterer zerebraler Strukturen unter transkutaner Vagusnervstimulation. 53:104–111.

28. Farmer AD et al. (2021) International Consensus Based Review and Recommendations for Minimum Reporting Standards in Research on Transcutaneous Vagus Nerve Stimulation (Version 2020). Frontiers in Human Neuroscience 14.

29. Ferstl M, Kuhnel A, Klaus J, Lin WM, Kroemer NB (2024) Non-invasive vagus nerve stimulation conditions increased invigoration and wanting in depression. Compr Psychiatry 132:152488.

30. Ferstl M, Teckentrup V, Lin WM, Kräutlein F, Kühnel A, Klaus J, Walter M, Kroemer NB (2022) Non-invasive vagus nerve stimulation boosts mood recovery after effort exertion. Psychol Med 52:3029–3039.

31. Frangos E, Ellrich J, Komisaruk BR (2015) Non-invasive Access to the Vagus Nerve Central Projections via Electrical Stimulation of the External Ear: fMRI Evidence in Humans. Brain Stimul 8:624–636.

32. Gerges ANH, Williams EER, Hillier S, Uy J, Hamilton T, Chamberlain S, Hordacre B Clinical application of transcutaneous auricular vagus nerve stimulation: a scoping review. Disability and Rehabilitation:1–31.

33. Giraudier M, Ventura-Bort C, Burger AM, Claes N, D’Agostini M, Fischer R, Franssen M, Kaess M, Koenig J, Liepelt R, Nieuwenhuis S, Sommer A, Usichenko T, Van Diest I, von Leupoldt A, Warren CM, Weymar M (2022) Evidence for a modulating effect of transcutaneous auricular vagus nerve stimulation (taVNS) on salivary alpha-amylase as indirect noradrenergic marker: A pooled mega-analysis. Brain Stimul 15:1378–1388.

34. Harrer M, Cuijpers P, Furukawa T, Ebert D (2021) Doing meta-analysis with R: A hands-on guide: Chapman and Hall/CRC.

35. Hedges LV, Olkin I (1984) Nonparametric estimators of effect size in meta-analysis. Psychological Bulletin 96:573–580.

36. Hein E, Nowak M, Kiess O, Biermann T, Bayerlein K, Kornhuber J, Kraus T (2013) Auricular transcutaneous electrical nerve stimulation in depressed patients: a randomized controlled pilot study. J Neural Transm (Vienna) 120:821–827.

37. Higgins JPT, Thompson SG, Spiegelhalter DJ (2008) A Re-Evaluation of Random-Effects Meta-Analysis. Journal of the Royal Statistical Society Series A: Statistics in Society 172:137–159.

38. Hulsey DR, Riley JR, Loerwald KW, Rennaker RL, Kilgard MP, Hays SA (2017) Parametric characterization of neural activity in the locus coeruleus in response to vagus nerve stimulation. Experimental Neurology 289:21–30.

39. Janner H, Klausenitz C, Gürtler N, Hahnenkamp K, Usichenko TI (2018) Effects of Electrical Transcutaneous Vagus Nerve Stimulation on the Perceived Intensity of Repetitive Painful Heat Stimuli: A Blinded Placebo- and Sham-Controlled Randomized Crossover Investigation. Anesthesia & Analgesia 126.

40. Jigo M, Carmel JB, Wang Q, Rodenkirch C (2024) Transcutaneous cervical vagus nerve stimulation improves sensory performance in humans: a randomized controlled crossover pilot study. Sci Rep 14:3975.

41. Johnson RL, Wilson CG (2018) A review of vagus nerve stimulation as a therapeutic intervention. J Inflamm Res 11:203–213.

42. Kemp J, Després O, Pebayle T, Dufour A (2014) Age-related decrease in sensitivity to electrical stimulation is unrelated to skin conductance: An evoked potentials study. Clinical Neurophysiology 125:602–607.

43. Keute M, Demirezen M, Graf A, Mueller NG, Zaehle T (2019) No modulation of pupil size and event-related pupil response by transcutaneous auricular vagus nerve stimulation (taVNS). Sci Rep-Uk 9:11452.

44. Kiyokawa J, Yamaguchi K, Okada R, Maehara T, Akita K (2014) Origin, course and distribution of the nerves to the posterosuperior wall of the external acoustic meatus. Anatomical Science International 89:238–245.

45. Konjusha A, Yu S, Mückschel M, Colzato L, Ziemssen T, Beste C (2023) Auricular Transcutaneous Vagus Nerve Stimulation Specifically Enhances Working Memory Gate Closing Mechanism: A System Neurophysiological Study. J Neurosci 43:4709–4724.

46. Kühnel A, Teckentrup V, Neuser MP, Huys QJM, Burrasch C, Walter M, Kroemer NB (2020) Stimulation of the vagus nerve reduces learning in a go/no-go reinforcement learning task. Eur Neuropsychopharmacol 35:17–29.

47. Lampros M, Vlachos N, Zigouris A, Voulgaris S, Alexiou GA (2021) Transcutaneous Vagus Nerve Stimulation (t-VNS) and epilepsy: A systematic review of the literature. Seizure 91:40–48.

48. Liu Y, Gelman A, Zheng T (2015) Simulation-efficient shortest probability intervals. Statistics and Computing 25:809–819.

49. Lloyd B, Wurm F, de Kleijn R, Nieuwenhuis S (2023) Short-term transcutaneous vagus nerve stimulation increases pupil size but does not affect EEG alpha power: A replication of Sharon et al. (2021, Journal of Neuroscience). Brain Stimulation: Basic, Translational, and Clinical Research in Neuromodulation 16:1001–1008.

50. Ludwig M, Pereira C, Keute M, Düzel E, Betts MJ, Hämmerer D (2024) Evaluating phasic transcutaneous vagus nerve stimulation (taVNS) with pupil dilation: the importance of stimulation intensity and sensory perception. bioRxiv:2024.2007.2027.605407.

51. Mathôt S (2018) Pupillometry: Psychology, Physiology, and Function. J Cogn 1:16. Mridha Z, de Gee JW, Shi Y, Alkashgari R, Williams J, Suminski A, Ward MP, Zhang W, McGinley MJ (2021) Graded recruitment of pupil-linked neuromodulation by parametric stimulation of the vagus nerve. Nature Communications 12:1539.

52. Neuser MP, Teckentrup V, Kühnel A, Hallschmid M, Walter M, Kroemer NB (2020) Vagus nerve stimulation boosts the drive to work for rewards. Nat Commun 11:3555.

53. Nieuwenhuis S, De Geus EJ, Aston-Jones G (2011) The anatomical and functional relationship between the P3 and autonomic components of the orienting response. Psychophysiology 48:162–175.

54. Omer S, Firas F, Yuval N (2021) Transcutaneous Vagus Nerve Stimulation in Humans Induces Pupil Dilation and Attenuates Alpha Oscillations. The Journal of Neuroscience 41:320.

55. Pandža NB, Phillips I, Karuzis VP, O’Rourke P, Kuchinsky SE (2020) Neurostimulation and Pupillometry: New Directions for Learning and Research in Applied Linguistics. Annual Review of Applied Linguistics 40:56–77.

56. Pike AC, Robinson OJ (2022) Reinforcement Learning in Patients With Mood and Anxiety Disorders vs Control Individuals: A Systematic Review and Meta-analysis. JAMA Psychiatry 79:313–322.

57. Polak T, Markulin F, Ehlis AC, Langer JB, Ringel TM, Fallgatter AJ (2009) Far field potentials from brain stem after transcutaneous vagus nerve stimulation: optimization of stimulation and recording parameters. J Neural Transm (Vienna) 116:1237–1242.

58. Preuschoff K, ’t Hart BM, Einhauser W (2011) Pupil Dilation Signals Surprise: Evidence for Noradrenaline’s Role in Decision Making. Frontiers in Neuroscience 5.

59. Redgrave J, Day D, Leung H, Laud PJ, Ali A, Lindert R, Majid A (2018) Safety and tolerability of Transcutaneous Vagus Nerve stimulation in humans; a systematic review. Brain Stimul 11:1225–1238.

60. Reimer J, McGinley MJ, Liu Y, Rodenkirch C, Wang Q, McCormick DA, Tolias AS (2016) Pupil fluctuations track rapid changes in adrenergic and cholinergic activity in cortex. Nature Communications 7:13289.

61. Rong P, Liu J, Wang L, Liu R, Fang J, Zhao J, Zhao Y, Wang H, Vangel M, Sun S, Ben H, Park J, Li S, Meng H, Zhu B, Kong J (2016) Effect of transcutaneous auricular vagus nerve stimulation on major depressive disorder: A nonrandomized controlled pilot study. J Affect Disord 195:172–179.

62. Röver C (2020) Bayesian Random-Effects Meta-Analysis Using the bayesmeta R Package. Journal of Statistical Software 93:1–51.

63. Röver C, Friede T (2023) Using the bayesmeta R package for Bayesian random-effects meta-regression. Comput Meth Prog Bio 229.

64. Sharon O, Fahoum F, Nir Y (2021) Transcutaneous Vagus Nerve Stimulation in Humans Induces Pupil Dilation and Attenuates Alpha Oscillations. J Neurosci 41:320–330.

65. Skora L, Marzecová A, Jocham G (2024) Tonic and phasic transcutaneous auricular vagus nerve stimulation (taVNS) both evoke rapid and transient pupil dilation. Brain Stimulation: Basic, Translational, and Clinical Research in Neuromodulation 17:233–244.

66. Sterne JAC et al. (2019) RoB 2: a revised tool for assessing risk of bias in randomised trials. Bmj 366:l4898.

67. Szeska C, Richter J, Wendt J, Weymar M, Hamm AO (2020) Promoting long-term inhibition of human fear responses by non-invasive transcutaneous vagus nerve stimulation during extinction training. Sci Rep 10:1529.

68. Tarasenko A, Guazzotti S, Minot T, Oganesyan M, Vysokov N (2022) Determination of the Effects of Transcutaneous Auricular Vagus Nerve Stimulation on the Heart Rate Variability Using a Machine Learning Pipeline. Bioelectricity 4:168–177.

69. Teckentrup V, Kroemer NB (2023) Mechanisms for Survival: Vagal Control of Goal-Directed Behavior. Trends Cogn Sci.

70. Teckentrup V, Neubert S, Santiago JCP, Hallschmid M, Walter M, Kroemer NB (2020) Non-invasive stimulation of vagal afferents reduces gastric frequency. Brain Stimul 13:470–473.

71. Teckentrup V, Krylova M, Jamalabadi H, Neubert S, Neuser MP, Hartig R, Fallgatter AJ, Walter M, Kroemer NB (2021) Brain signaling dynamics after vagus nerve stimulation. NeuroImage 245:118679.

72. Tekdemir I, Aslan A, Elhan A (1998) A clinico-anatomic study of the auricular branch of the vagus nerve and Arnold’s ear-cough reflex. Surg Radiol Anat 20:253–257.

73. Urbin MA, Lafe CW, Simpson TW, Wittenberg GF, Chandrasekaran B, Weber DJ (2021) Electrical stimulation of the external ear acutely activates noradrenergic mechanisms in humans. Brain Stimulation 14:990–1001.

74. Vespa S, Stumpp L, Liberati G, Delbeke J, Nonclercq A, Mouraux A, El Tahry R (2022). Characterization of vagus nerve stimulation-induced pupillary responses in epileptic patients. Brain Stimul 15:1498–1507.

75. Viechtbauer W (2010) Conducting meta-analyses in R with the metafor package. Journal of statistical software 36:1–48.

76. Villani V, Finotti G, Di Lernia D, Tsakiris M, Azevedo RT (2022) Event-related transcutaneous vagus nerve stimulation modulates behaviour and pupillary responses during an auditory oddball task. Psychoneuroendocrinology 140:105719.

77. Wang C-A, Munoz DP (2015) A circuit for pupil orienting responses: implications for cognitive modulation of pupil size. Current Opinion in Neurobiology 33:134–140.

78. Wang Y, Zhan G, Cai Z, Jiao B, Zhao Y, Li S, Luo A (2021) Vagus nerve stimulation in brain diseases: Therapeutic applications and biological mechanisms. Neurosci Biobehav Rev 127:37–53.

79. Warren CM, Tona KD, Ouwerkerk L, van Paridon J, Poletiek F, van Steenbergen H, Bosch JA, Nieuwenhuis S (2019) The neuromodulatory and hormonal effects of transcutaneous vagus nerve stimulation as evidenced by salivary alpha amylase, salivary cortisol, pupil diameter, and the P3 event-related potential. Brain Stimul 12:635–642

80. Weber I, Niehaus H, Krause K, Molitor L, Peper M, Schmidt L, Hakel L, Timmermann L, Menzler K, Knake S, Oehrn CR (2021) Trust your gut: vagal nerve stimulation in humans improves reinforcement learning. Brain Commun 3:fcab039.

81. Wienke C, Grueschow M, Haghikia A, Zaehle T (2023) Phasic, Event-Related Transcutaneous Auricular Vagus Nerve Stimulation Modifies Behavioral, Pupillary, and Low-Frequency Oscillatory Power Responses. The Journal of Neuroscience 43:6306.

82. Wolf V, Kühnel A, Teckentrup V, Koenig J, Kroemer NB (2021) Does transcutaneous auricular vagus nerve stimulation affect vagally mediated heart rate variability? A living and interactive Bayesian meta-analysis. Psychophysiology 58:e13933.

83. Yakunina N, Kim SS, Nam EC (2017) Optimization of Transcutaneous Vagus Nerve Stimulation Using Functional MRI. Neuromodulation 20:290–300.

84. Yakunina N, Kim SS, Nam E-C (2018) BOLD fMRI effects of transcutaneous vagus nerve stimulation in patients with chronic tinnitus. PLOS ONE 13:e0207281.

85. Yang J, Phi JH (2019) The Present and Future of Vagus Nerve Stimulation. J Korean Neurosurg Soc 62:344–352.

86. Yap JYY, Keatch C, Lambert E, Woods W, Stoddart PR, Kameneva T (2020) Critical Review of Transcutaneous Vagus Nerve Stimulation: Challenges for Translation to Clinical. Practice. Frontiers in Neuroscience 14.

