## Supporting Information for "Does transcutaneous vagus nerve stimulation alter pupil dilation? A living Bayesian meta-analysis"

Venusberg Campus 1, 53127 Bonn, Germany

### Search strategies

The systematic literature search was performed on 17/10/2023 in the databases Pubmed and Web of Science. The following keywords were used for the search: (VNS OR “vagus nerve stimulation” OR “vagal nerve stimulation” OR tVNS OR taVNS OR atVNS OR “non-invasive vagus nerve stimulation” OR “transcutaneous vagus nerve stimulation” OR “transcutaneous auricular vagus nerve stimulation” OR ”transcutaneous vagal nerve stimulation” OR “non-invasive vagal nerve stimulation” OR “transcutaneous auricular vagal nerve stimulation” OR “vagus” OR “vagus nerve” OR “vagal nerve”) AND (”eye tracking” OR pupil OR “pupil dilation” OR “pupil size” OR “pupil diameter” OR pupillometry OR “pupillary light reflex” OR “pupillography”). Relevant papers from both databases were combined and duplicates were removed. In addition to the papers identified through the systematic database search, three papers were manually added after identification from preprint servers or notification by the authors. After screening abstracts, 20 papers were assessed. From these 20 papers, 3 papers had to be excluded (one did not include sham stimulation as control (McHaney et al., 2023), one used percutaneous VNS (Treiber et al., 2024) and one did not record pupil dilation during tVNS, but only after the stimulation period (Zhu et al., 2022)), leaving 17 papers that were relevant for the meta-analysis (Figure 1). For some papers (Keute et al., 2019; Warren et al., 2019; Burger et al., 2020; D’Agostini et al., 2021; Urbin et al., 2021; D'Agostini et al., 2022; D'Agostini et al., 2023; Lloyd et al., 2023) that did not report summary data on pupil dilation, we extracted information from their figures using *WebPlotDigitizer* (Version 4.6, https://automeris.io/WebPlotDigitizer).

**tVNS Stimulation**

taVNS stimulation is applied using various protocols with two broad categories emerging. First, in line with most clinical applications (Gerges et al.), the vagus nerve is stimulated continuously over several minutes to hours. During this phase, stimulation can either be biphasic waves that are often applied with ~30-second ON/ OFF protocols (Warren et al., 2019; Burger et al., 2020; D'Agostini et al., 2021; Skora et al., 2024), or monophasic waves delivered throughout the whole stimulation duration (Keute et al., 2019; Borges et al., 2021; Capone et al., 2021; D'Agostini et al., 2022). Second, inspired by animal research, the vagus nerve can be stimulated with a pulsed protocol that entails brief stimulation periods of a few seconds (0.6s (Urbin et al., 2021) - 5s (D'Agostini et al., 2023)) as would be expected by bodily feedback signals. For coding the studies, stimulation periods shorter or equal to 5s were considered as pulsed stimulation. Continuous stimulation protocols had at least 30s of uninterrupted stimulation.

In addition to the two stimulation protocol types, taVNS studies vary in the applied frequency, pulse width, and intensity. In the studies included in this meta-analysis, the applied frequency ranges between 20 Hz (Capone et al., 2021) and 300 Hz (Urbin et al., 2021), while pulse widths vary from 200 µs (Keute et al., 2019; Wienke et al., 2023) to 400 µs (D'Agostini et al., 2023). Notably, the stimulation intensity is another point of divergence among studies, with 6 studies (2 continuous taVNS, 4 pulsed taVNS) determining it based on participants' perceptual threshold (Urbin et al., 2021). reach it leading to different sensations for sham and tVNS (D'Agostini et al., 2021). Another 8 studies (6 continuous taVNS, 3 pulsed taVNS) used fixed intensity for all participants with no difference between the stimulation intensity applied during taVNS and sham (e.g., (Warren et al., 2019)) sometimes reducing individual intensities when pain was reported (Wienke et al., 2023; Ludwig et al., 2024). One study (pulsed and continuous tVNS) used a pain threshold but did not calibrate for each condition (sham vs. tVNS) separately but just for the first participants received (Skora et al., 2024) so sensation ratings are not necessarily matched. Further, one study stimulation intensity was calibrated based on ratings but with a relatively low maximum stimulation amplitude and 95% of the participants did. The study from D'Agostini and colleagues (2023) included effects both from fixed stimulation intensities and sensation-matched intensities although ratings after stimulation still differed between sham and tVNS. After establishing the desired threshold, one study further explored different amplitude multiples of the individually calibrated intensity (Urbin et al., 2021).

**Pupillometry**

Pupil dilation is measured using eye tracking and different measurement devices were used across the included studies. Specifically, 3 papers used SensoMotoric Instrument eye-tracking glasses (Warren et al., 2019; Borges et al., 2021; D’Agostini et al., 2021), 3 used a Tobii T120 eye-tracker (Burger et al., 2020; Villani et al., 2022; Lloyd et al., 2023), and one used a Sirius 3D rotating camera topographic system (Capone et al., 2021). Pupil size was measured with an SR-Research Eyelink 1000 in 8 studies (Keute et al., 2019; Ludwig et al., 2021; Sharon et al., 2021; Urbin et al., 2021; D'Agostini et al., 2022; D'Agostini et al., 2023; Wienke et al., 2023; Skora et al., 2024).

Pupil size can be recorded either monocularly (11 studies) or binocularly (3 studies; 5 studies did not specify the side of recording) with subsequent averaging between the left and right eye. For continuous tVNS, the majority of papers analyzed changes in pupil size by averaging across the complete stimulation period. Continuous analyses capture changes in pupil size over a longer time and are expressed as baseline-corrected dilation in 12 studies (7 continuous taVNS, 4 pulsed taVNS, 1 both; e.g., (Keute et al., 2019) or non-corrected pupil size in 5 studies (3 continuous taVNS, 2 pulsed taVNS; e.g., (Capone et al., 2021)). If studies are using experimental tasks (e.g. an oddball task), another common analysis method for pupil size is the determination of event-related pupil dilation (ERPD). ERPDs for task events are the maximum post-event deviation from the pre-event pupil size baseline for each trial of the experimental task (see (D'Agostini et al., 2022; Villani et al., 2022) for an example). When this is combined with tVNS, the outcome is a tVNS-induced change in the task-event related pupil dilation. For pulsed tVNS, pupil dilations are measured for short (a few seconds long) post-stimulation epochs and the maximal dilation per trial is determined in comparison to a pre-stimulus baseline comparable to the approach for task event-related pupil dilations (see e.g., (D'Agostini et al., 2023)). Skora and colleagues also used this event-related phasic analysis approach for their continuous taVNS stimulation (Skora et al., 2024). Comparable to ERPDs, they analyzed pupil response in the short epochs (~5s) starting from the stimulation onset of each 30s ON phase and baseline-corrected using the last seconds of the previous 30s OFF period (Skora et al., 2024).

**Coding procedure**

For all screened papers, we (IP and LT) coded the type of stimulation (pulsed, continuous), stimulation side, part of the ear stimulated, frequency, pulse width, stimulation (pulse) time (starting from 0.6 s – 5s for the pulsed protocols and either 30s or the complete stimulation time if stimulation was monophasic for continuous stimulation), stimulation duration during the whole experiment and amplitude of the delivered stimulation, the type of eye-tracking recording (monocular, binocular), eye-tracking device and duration of the recording. We also reported whether the recording was done during a task and whether the outcome was a phasic pupil dilation in response to an event or an average across a longer time.

**Risk of bias**

The risk of bias was assessed using the open-access revised Cochrane risk of bias tool for randomized trials *(RoB2*, (Sterne et al., 2019)*).* Two of the authors (IP and LK) conducted the assessment independently. All studies were categorized as low risk concerning randomization. One study (Capone et al., 2021) provided no information regarding missing data leading to the categorization of some risks in this domain and for successful blinding. One study (D'Agostini et al., 2021) is categorized as high risk in domain 2 (blinding) because blinding could not be guaranteed according to a post stimulation assessment of the participants. We categorized ten studies (Keute et al., 2019; Warren et al., 2019; Burger et al., 2020; D'Agostini et al., 2021; D'Agostini et al., 2023; Wienke et al., 2023; Ludwig et al., 2024; Skora et al., 2024) as high risk regarding the validity of the measurement because the stimulation intensities were not succesfully matched for tVNS and sham based on the self-reported sensation and pupil size is also affected by sensory stimulation. Of the studies pre-registering their analysis, only three followed it as described (D'Agostini et al., 2023; Lloyd et al., 2023; Skora et al., 2024) whereas the others did not preregister their analysis, diverged from the preregistration, or had the risk of not reporting all performed analyses.

**Design of the Shiny app**

To build the Shiny app, we adapted a previously developed app by our group (Wolf et al., 2021). Analogous to the previous living meta-analysis using taVNS-induced changes in heart rate variability as the outcome, the Shiny app allows the user to change the inclusion criteria for the selection of studies into the meta-analysis. For example, users may select the percentage of women present in the sample, the mean age of the sample, the location of electrode placement (i.e. cymba concha, tragus, ear canal), and the publication year(s). Furthermore, it allows the selection of different parameters, such as the type of stimulation (continuous, phasic), the duration of stimulation pulses, the total duration of stimulation during the experiment, the duration of the electrophysiological recording, the recorded eye (left, right, or both), the presence of a task during stimulation, and the pupil size measurement (i.e., tonic increase or event-related pupil dilation).

Ludwig M, Wienke C, Betts MJ, Zaehle T, Hämmerer D (2021) Current challenges in reliably targeting the noradrenergic locus coeruleus using transcutaneous auricular vagus nerve stimulation (taVNS). Autonomic Neuroscience 236:102900.

Ludwig M, Pereira C, Keute M, Düzel E, Betts MJ, Hämmerer D (2024) Evaluating phasic transcutaneous vagus nerve stimulation (taVNS) with pupil dilation: the importance of stimulation intensity and sensory perception. bioRxiv:2024.2007.2027.605407.

McHaney JR, Schuerman WL, Leonard MK, Chandrasekaran B (2023) Transcutaneous Auricular Vagus Nerve Stimulation Modulates Performance but Not Pupil Size During Nonnative Speech Category Learning. J Speech Lang Hear Res 66:3825-3843.

Sharon O, Fahoum F, Nir Y (2021) Transcutaneous Vagus Nerve Stimulation in Humans Induces Pupil Dilation and Attenuates Alpha Oscillations. J Neurosci 41:320-330.
Skora L, Marzecová A, Jocham G (2024) Tonic and phasic transcutaneous auricular vagus nerve stimulation (taVNS) both evoke rapid and transient pupil dilation. Brain Stimulation: Basic, Translational, and Clinical Research in Neuromodulation 17:233-244.
Sterne JAC et al. (2019) RoB 2: a revised tool for assessing risk of bias in randomised trials. Bmj 366:l4898.
Treiber MC, Grünberger J, Vyssoki B, Szeles JC, Kaniusas E, Kampusch S, Stöhr H, Walter H, Lesch OM, König D, Kraus C (2024) Pupillary response to percutaneous auricular vagus nerve stimulation in alcohol withdrawal syndrome: A pilot trial. Alcohol 114:61-68.
Urbin MA, Lafe CW, Simpson TW, Wittenberg GF, Chandrasekaran B, Weber DJ (2021) Electrical stimulation of the external ear acutely activates noradrenergic mechanisms in humans. Brain Stimulation 14:990-1001.
Villani V, Finotti G, Di Lernia D, Tsakiris M, Azevedo RT (2022) Event-related transcutaneous vagus nerve stimulation modulates behaviour and pupillary responses during an auditory oddball task. Psychoneuroendocrinology 140:105719.
Warren CM, Tona KD, Ouwerkerk L, van Paridon J, Poletiek F, van Steenbergen H, Bosch JA, Nieuwenhuis S (2019) The neuromodulatory and hormonal effects of transcutaneous vagus nerve stimulation as evidenced by salivary alpha amylase, salivary cortisol, pupil diameter, and the P3 event-related potential. Brain Stimul 12:635-642.
Wienke C, Grueschow M, Haghikia A, Zaehle T (2023) Phasic, Event-Related Transcutaneous Auricular Vagus Nerve Stimulation Modifies Behavioral, Pupillary, and Low-Frequency Oscillatory Power Responses. The Journal of Neuroscience 43:6306.
Wolf V, Kühnel A, Teckentrup V, Koenig J, Kroemer NB (2021) Does transcutaneous auricular vagus nerve stimulation affect vagally mediated heart rate variability? A living and interactive Bayesian meta-analysis. Psychophysiology 58:e13933.
Zhu S, Qing Y, Zhang Y, Zhang X, Ding F, Zhang R, Yao S, Kendrick KM, Zhao W (2022) Transcutaneous auricular vagus nerve stimulation increases eye-gaze on salient facial features and oxytocin release. Psychophysiology 59:e14107.
